# Shape coding in occipito-temporal cortex relies on object silhouette, curvature and medial-axis

**DOI:** 10.1101/814251

**Authors:** Paolo Papale, Andrea Leo, Giacomo Handjaras, Luca Cecchetti, Pietro Pietrini, Emiliano Ricciardi

## Abstract

Object recognition relies on different transformations of the retinal input, carried out by the visual system, that range from local contrast to object shape and category. While some of those transformations are thought to occur at specific stages of the visual hierarchy, the features they represent are correlated (e.g., object shape and identity) and selectivity for the same feature overlaps in many brain regions. This may be explained either by collinearity across representations, or may instead reflect the coding of multiple dimensions by the same cortical population. Moreover, orthogonal and shared components may differently impact on distinctive stages of the visual hierarchy. We recorded functional MRI (fMRI) activity while participants passively attended to object images and employed a statistical approach that partitioned orthogonal and shared object representations to reveal their relative impact on brain processing. Orthogonal shape representations (silhouette, curvature and medial-axis) independently explained distinct and overlapping clusters of selectivity in occitotemporal (OTC) and parietal cortex. Moreover, we show that the relevance of shared representations linearly increases moving from posterior to anterior regions. These results indicate that the visual cortex encodes shared relations between different features in a topographic fashion and that object shape is encoded along different dimensions, each representing orthogonal features.

**New & Noteworthy:** There are several possible ways of characterizing the shape of an object. Which shape description better describes our brain responses while we passively perceive objects? Here, we employed three competing shape models to explain brain representations when viewing real objects. We found that object shape is encoded in a multi-dimensional fashion and thus defined by the interaction of multiple features.

## Introduction

Objects are often defined by their shape. However, from a theoretical perspective, the concept of shape may appear quite elusive, since the shape of an object could be defined in several possible ways. Consequently, many computational models of object shape can be constructed: closed shapes can be easily and reliably generated by combining simple elements (e.g., *geons* or medial axes), or by connecting few salient points with acute curvature, or by modulating radial frequency. Hence, we can ask which of these different descriptions is encoded in our brain and reflects the way we perceive and represent shapes. Here, we show that shape coding in the brain relies on multiple dimensions.

Indeed, all these different representations are sensitive to specific aspects of a shape and produce different ways of encoding the same object. Consider Figure 1. A silhouette descriptor (Figure 1B and 2A, red), consisting of a simple stimulus vectorization, would vary mostly depending on the global layout of a shape, but would be quite insensitive to small perturbations of the outline. Conversely, the curvature descriptor (Figure 1B and 2A, blue) would be unaffected by rotations or large transformation of a shape, but would be highly sensitive to the number of acute points on the outline. Of note, these different representations can either diverge independently to each other, as in the horizontal and vertical shapes, or can covary together, as for the shapes placed on the diagonal. Evidence from previous studies suggests that visual dimensions in natural vision are indeed highly correlated (Figure 2D; Bracci and Op de Beeck 2016; Kay 2011; Papale et al. 2019). Thus, addressing the extent to which brain regions represent different dimensions has so far proven challenging: how can we disentangle the role of different shape properties if they likely covary together?

**Figure 1.**
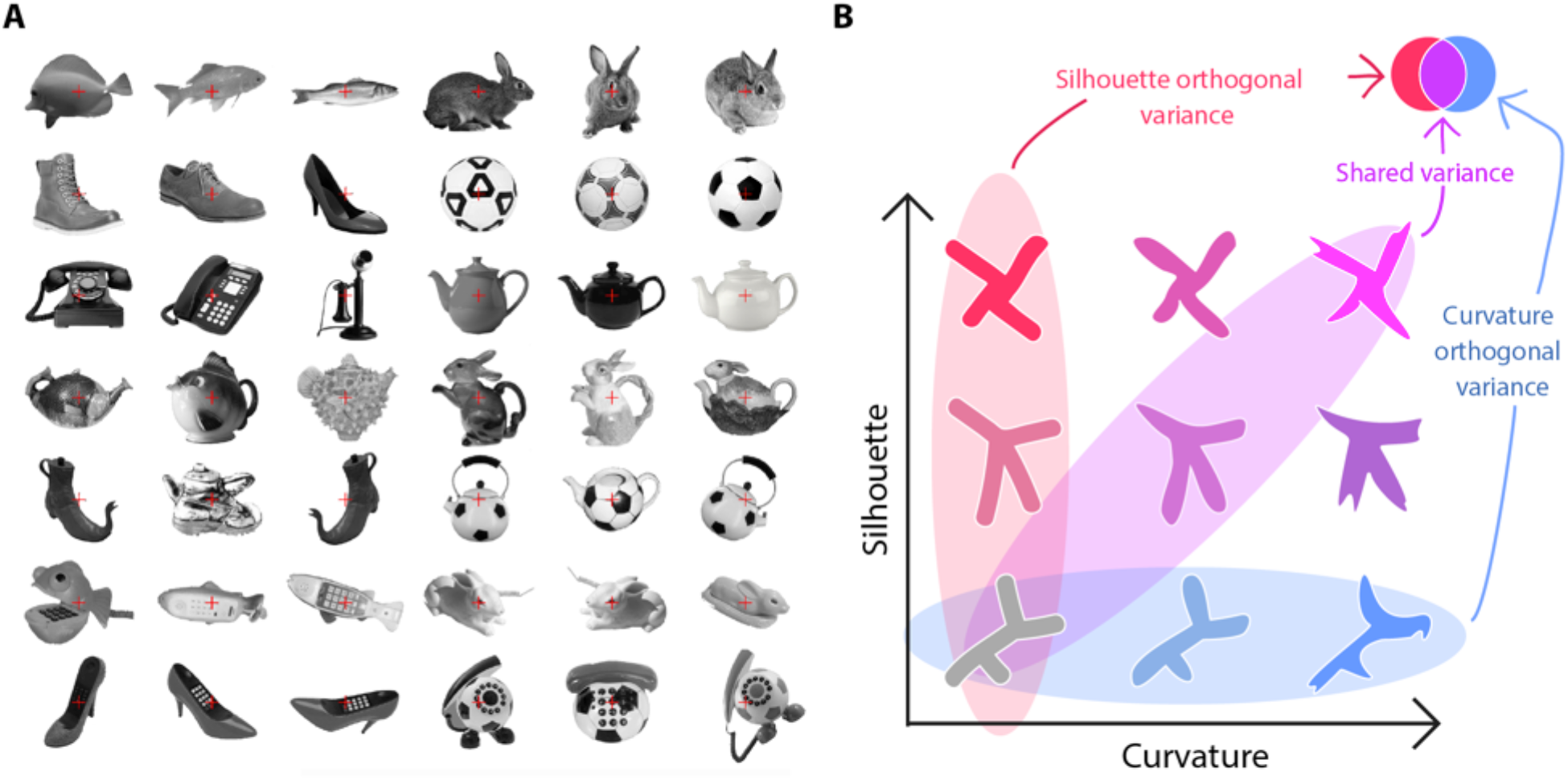
How does our brain encode object shape? A) The full stimulus set used in our study. As evident, some of the images depict daily, familiar objects, while some of them show unambiguous hybrids, combining two familiar objects. B) Different features capture specific aspects of object shape. For instance, silhouette and curvature descriptions of the same shapes may be orthogonal to each other (red- and blue-shaded areas) or vary in a linear fashion (purple-shaded area). Thus, our brain may represent object shape by extracting one specific and more reliable feature, by focusing on shared representations across multiple features, or even encoding the orthogonal components of different features.

**Figure 2.**
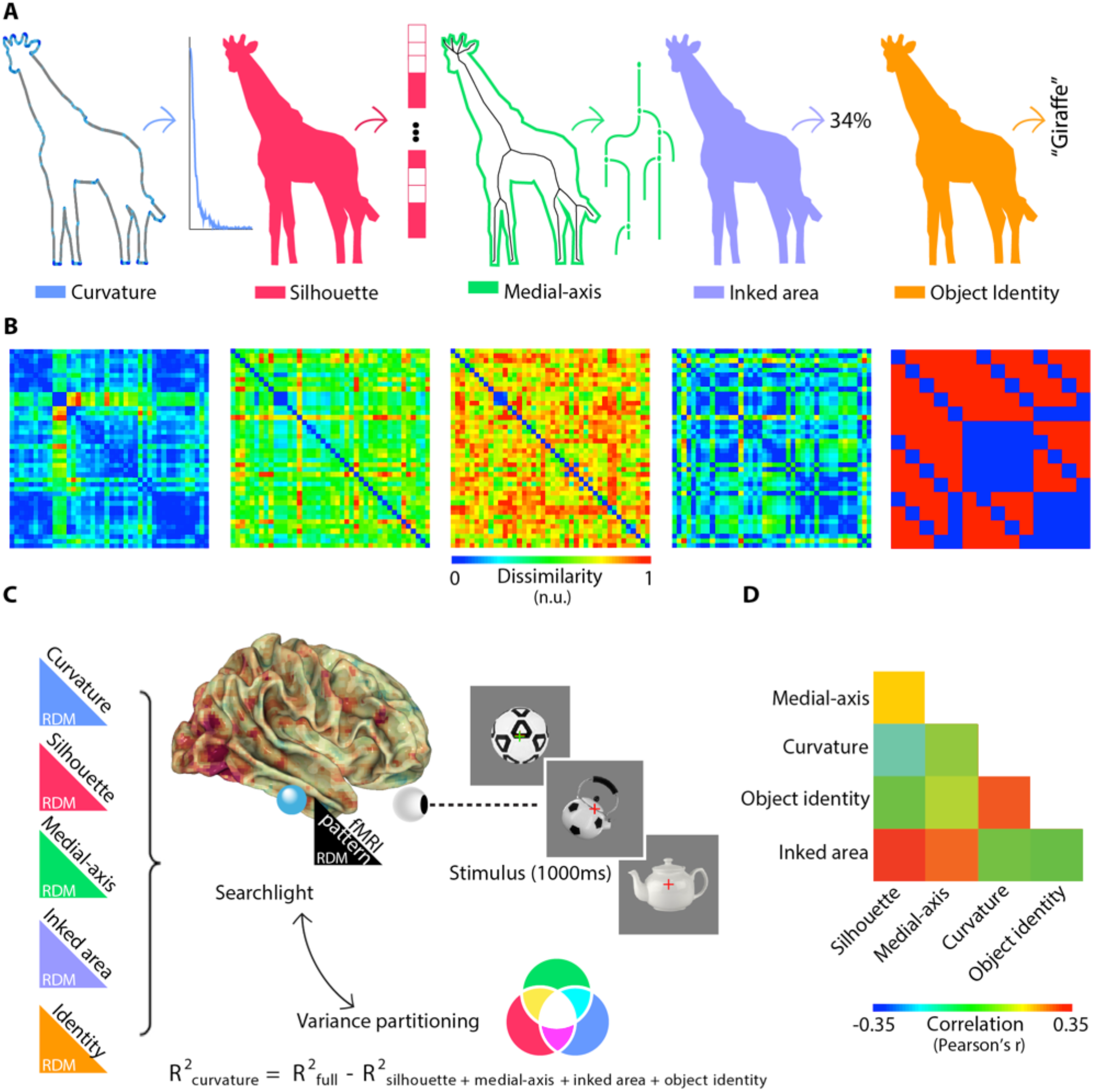
Schematic of the shape models and experiment. A) Five different object representations are employed: three shape models and two further descriptions. From left: silhouette, medial axis, curvature, inked area (low-level) and object identity (high-level). B) Representational dissimilarity matrices (RDMs) of each model: they represent all the possible pairwise distances between the stimuli. C) Methodological pipeline. Brain responses were recorded while subjects maintained fixation on a colored fixation cross, paying attention to color switching between red and green. Orthogonal to the task, we presented 42 grayscale pictures of real objects, for 1s each. Activity patterns were used to test the association between the five model RDMs and each brain activity RDM, computed combining a searchlight procedure with a variance partitioning analysis. D) Similarity between the five model RDMs. As expected, the five representations are correlated. However, the variance partitioning approach controls for the effect of model collinearity.

To answer these questions, we recorded functional MRI (fMRI) activity while participants passively viewed object pictures (Figure 1A). We employed a statistical approach that partitions orthogonal and shared shape representations and reveals their relative impact on brain processing, while controlling, at the same time, for low- and high-level features (Figure 2C; Greene et al. 2016; Groen et al. 2012; Groen et al. 2018; Hebart et al. 2018; Lescroart et al. 2015; Ramakrishnan et al. 2014).

Since a recent behavioral work demonstrates that multiple dimensions are necessary to model behavioral shape similarity (Morgenstern et al. 2020), we expected multiple shape properties to be independently represented in the human visual cortex. At the same time, we explored the way in which the brain exploits shared information between different visual dimensions. In line with recent neurophysiological work (Hong et al. 2016), we expected to observe a linear decrease of the role of orthogonal representations along the visual hierarchy.

In our approach, we tested three competing shape models. A first description was computed by extracting the silhouette. The link between shape silhouette and representations in the occipito-temporal cortex (OTC) has been extensively investigated in neuroimaging studies (Bracci and Op de Beeck 2016; Kaiser et al. 2016; Khaligh-Razavi and Kriegeskorte 2014; Proklova et al. 2016). Second, a skeletal representation of each stimulus was extracted by performing the medial axis transform (Blum 1973). The spike rate of inferotemporal (IT) neurons in monkey are sensitive to the medial-axis of objects (Hung et al. 2012), which also captures behavioral ratings of shape similarity (Ayzenberg et al. 2019a; Ayzenberg and Lourenco 2019; Lowet et al. 2018) and whose spatiotemporal association with brain activity in humans has been described in several neuroimaging studies (Ayzenberg et al. 2019b; Handjaras et al. 2017; Leeds et al. 2013; Lescroart and Biederman 2013; Papale et al. 2019). A third description was obtained by computing the curvature distribution for each object contour. It has been showed that V4 neurons in monkey are selective to a specific degree of curvature (Cadieu et al. 2007; Carlson et al. 2011; Connor et al. 2007). Moreover, the pivotal role of contour curvature in object perception has been extensively demonstrated both by behavioral (Elder and Velisavljevic 2009; Lawrence et al. 2016; Long et al. 2017; Wolfe et al. 1992) and neuroimaging studies (Caldara et al. 2006; Long et al. 2018; Vernon et al. 2016; Yue et al. 2014). In addition, we introduced two additional models: the inked area (in pixels) of each stimulus as a low-level representation, and object identity, to account for high-level information (Khaligh-Razavi and Kriegeskorte 2014; Kriegeskorte et al. 2008). Of note, all these models differ in their complexity: the silhouette and inked area return a low-level description of which parts and what extent of the visual field (and thus of the retinotopic cortex) are covered by the picture of an object, respectively. Instead, medial-axis, curvature and identity provide high-level, abstract representations, insensitive to viewpoint and local position.

As expected, we found both distinct and overlapping clusters of selectivity in OTC and in parietal regions independently explained by different shape representations (i.e., silhouette, curvature and medial-axis). Moreover, we showed that, while the prominence of retinotopic processing on abstract information shifts abruptly moving from the occipital to the temporal cortex, shared representations linearly increase from posterior to anterior regions along the visual hierarchy.

## Materials and Methods

### Subjects

Seventeen subjects were enrolled for the study. Two subjects participated as pilot subjects with a different version of the experimental protocol and their data were not used for the subsequent analyses; data from one subject who abruptly terminated the experiment were discarded. Fourteen subjects were further considered. The final sample comprised six females, age was 24 ± 3 years, all subjects were right-handed with normal or corrected-to-normal vision and were recruited among the students at the University of Pisa, Italy. Signed informed consent was acquired from all subjects and all the experimental procedures were performed according to the Declaration of Helsinki, under a protocol (1616/2003) approved by the Ethical Committee at the University of Pisa, Italy.

### Task

For this study, an event-related design was adopted. Stimuli consisted of 42 static images of grayscale unfamiliar and common objects (Figure 1A), presented against a fixed gray background, with a superimposed fixation cross (size: 2×2°), followed by a baseline condition characterized by a gray screen with a red fixation cross.

A set of stimuli was selected, consisting of 24 common (animate and inanimate) and 18 unfamiliar objects (500×500 pixels – 20° ×20°). The latter group represented existing objects that combine the function and the shape of two of the common objects (e.g., a fish-shaped teapot). Of note, a similar criterion has been employed for stimuli selection also in a recent study (Bracci et al. 2019). To build the final set of stimuli, pictures of existing objects were found on Internet, resized, normalized for luminance and root-mean-square contrast.

Stimuli were presented with the Presentation software (Neurobehavioral Systems, Albany, CA, USA) on MR-compatible goggles (VisuaStim, Resonance Technology Inc., CA, USA), with a LCD at the resolution of 800×600 pixels (32° ×24°). The study was organized in six runs, comprising 56 trials (in total: 8 repetitions for each stimulus) which consisted of 1000ms of stimulus presentation and 7000ms of inter-stimulus interval; each run started and ended with 15 seconds of rest, to estimate baseline levels of BOLD signal, and lasted 7:30 minutes. The total duration of the experiment, including anatomical scans, was about 55 minutes.

During the functional runs, subjects were asked to fixate the cross at the center of the screen. On selected trials, the cross changed its color from red to green, and subjects were asked to detect such changes by pressing a key on a MR-compatible keyboard with the index finger of their dominant hand. Order of trials was randomized across runs, and a different randomization schema was used for each participant.

### Functional MRI data acquisition

Data were acquired with a 3-Tesla GE Signa scanner (General Electric Inc., Milwaukee, WI, USA) equipped with an 8-channel phased-array coil. For functional images, a gradient-echo echo-planar imaging sequence (GE-EPI) was used, with TE = 40ms, TR = 2500ms, FA = 90°, 184 volumes, acquisition time 7’40”, including four additional dummy scans; image geometry parameters were: Field-Of-View 258×258mm, 128×128 in-plane matrix, voxel size 2.03×2.03×4mm, 37 axial slices for total brain coverage (z-axis extent = 148mm). To acquire detailed information of subject anatomy, a 3D Fast Spoiled Gradient Echo T1-weighted sequence was also acquired (TE = 3.18ms, TR = 8.16ms, FA = 12°, Field-Of-View 256×256mm, 256×256 matrix size, 1mm^3^ isotropic voxels, 256 axial slices, z-axis extent 256mm).

### Functional MRI data processing

Data preprocessing was carried out with AFNI (Cox 1996) and FSL 5.0 (Jenkinson et al. 2012). Preprocessing of functional data comprised slice timing correction with Fourier method (*3dTshift*), rigid-body motion correction using the first volume of the third run as reference (*3dvolreg*), spike removal (elimination of outliers in the functional time series, *3dDespike*), smoothing with a Gaussian filter (fixed FWHM 4 mm, *3dmerge*), and scaling of BOLD time series to percentage of the mean of each run (*3dTstat, 3dcalc*). Processing of anatomical images consisted of brain extraction (*bet*), segmentation for bias-field estimation and removal (*FAST, fslmaths*), linear (*FLIRT*) and nonlinear registration (*FNIRT*) to MNI152 standard space.

For each subject, data from the six concatenated runs (960 time points) were used for a GLM analysis (*3dDeconvolve*) with the responses for each stimulus – modeled with 1 seconds-long block functions convolved with a canonical HRF – as predictors of interest, and the six motion parameters plus polynomial trends up to 4^th^ order as predictors of no-interest.

Responses for individual stimuli were converted to MNI152 space by applying the transformation matrices estimated as explained above, and resampled to a resolution of 2×2×2mm.

### Computational models

Five different representations of the 42 stimuli were developed: three shape-based descriptions of interest and two further models. For each model, we obtained a stimulus-specific feature space, and pairwise dissimilarities between stimuli were computed to obtain a representational dissimilarity matrix (RDM). Before computing shape-related information, stimuli were binarized.

For the silhouette model, pairwise dissimilarity was computed using correlation distance (1 – Pearson’s r). For the medial-axis model, pairwise distance between skeletal representations was computed using the ShapeMatcher algorithm (http://www.cs.toronto.edu/~dmac/ShapeMatcher/; (Van Eede et al. 2006)). In sum, the ShapeMatcher algorithm builds the shock-graphs of each shape and then estimates their dissimilarity as the deformation required to match different objects (Sebastian et al. 2004). Curvature was computed as the chord-to-point distance (Monroy et al. 2011) in a 40-pixels window. Pairwise dissimilarity was computed using correlation distance between the histograms of curvature from each pair of stimuli. Finally, two further RDMs were built. For the inked-area, pairwise dissimilarity was computed as the absolute difference between the number of pixels covered by different objects. For identity, a binary representation was employed (Khaligh-Razavi and Kriegeskorte 2014; Kriegeskorte et al. 2008). Unfamiliar stimuli were considered as belonging to categories according to both their function and shape.

### Variance partitioning

A variance partitioning analysis (Lescroart et al. 2015) was performed to determine whether the three shape models in this study significantly explain unique components of the variance of brain representations (computed using Pearson’s correlation distance), as computed in 6 mm-radius spherical searchlights (Kriegeskorte et al. 2006). To this aim, explained variance coefficient (R^2^) was computed for each model RDM in independent linear regressions, and then all the different combinations of models were tested in further multiple linear regressions. The final statistic reporting the partial goodness of fit for unique and shared components was computed following the work by Nimon and colleagues (2008). For example, the unique variance explained by the curvature model in a specific searchlight was determined as the difference between the full-model R^2^ and the variance explained by the combination of all other models (i.e., R^2^ _curvature_ = R^2^_full_ – R^2^_silhouette + medial-axis + inked area + identity_). In the context of multiple linear regression, this approach is better known as ‘commonality analysis’ (Nimon and Oswald 2013), and its popularity is growing in neuroimaging (de Heer et al. 2017; Groen et al. 2018; Lescroart et al. 2015).

It should be noted that the so obtained orthogonal/unique partitions are not strictly uncorrelated from a statistical viewpoint (Creager 1971). However, referring to this procedure as *orthogonalization*, even though not fully accurate, yet makes its core philosophy clearer and easier to understand than the term *residualisation* that would be more statistically correct.

### Single-subject encoding

Correlation distance (1 – Pearson’s r) was used to compute the RDM of fMRI activity patterns in each searchlight and each subject. Only voxels pertaining to the cerebral cortex with a probability higher than 50% were included in the procedure (i.e., by applying a threshold over the Harvard-Oxford cortical probabilistic atlas). After applying variance partitioning, the obtained R^2^ for each component of unique and shared variance in each subject were z-scored, turned in the partial correlation coefficient (de Heer et al. 2017) and then assigned to the center of the searchlight, so obtaining a map for each subject and component. Results from four representative subjects are shown in Figures S1-4, while the similarity between the fittings of the full-model across subjects is shown in Figure S5.

### Group-level test

For each model, threshold free cluster enhancement (TFCE: Smith and Nichols 2009) was used to detect group-level clusters significantly explained by the corresponding unique variance component (5000 randomizations of the sign with 6mm variance smoothing, as implemented in FSL’s *randomise*: www.fmrib.ox.ac.uk/fsl/randomise). Statistical maps were then thresholded at one-tailed p < 0.05, corrected for multiple comparisons across gray matter voxels and finally a further arbitrary cluster size threshold of 10 voxels was applied (Figure 3).

**Figure 3.**
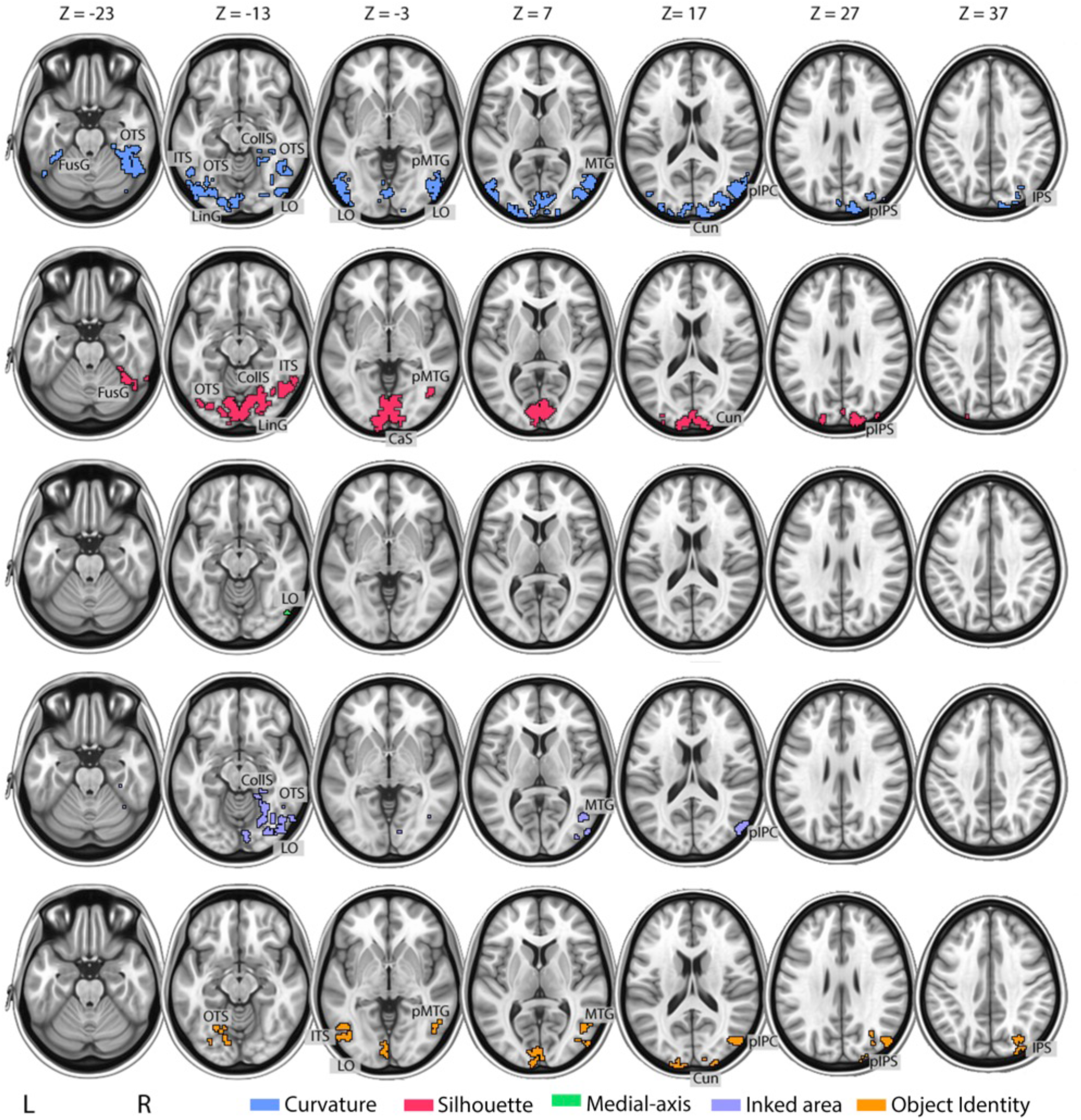
The human visual cortex encodes orthogonal shape representations. Group-level maps showing significant clusters of shape selectivity in OTC and in posterior dorsal regions (one-tailed p < 0.05, TFCE corrected). Each row of images depicts selectivity to orthogonal components of curvature (blue), silhouette (red), medial-axis (green), inked area (purple) and object identity (orange). CalcS: Calcarine sulcus; OTS: occipitotemporal sulcus; CollS: collateral sulcus; ITS: inferior temporal sulcus; FusG: fusiform gyrus; Cun: cuneus; MTG: middle temporal gyrus; IPS: intraparietal sulcus; IPC: intraparietal cortex.

### Orthogonality and complexity testing

To look for differences in how information is encoded in different brain regions, we introduced the orthogonality and complexity measures. Orthogonality was computed by dividing the group-averaged (between subjects) sum of variance explained uniquely by the five models with the group-averaged (between subjects) sum of variance explained by their shared components for each searchlight (Orthogonality = Σ R^2^ _unique components_ / Σ R2 _shared components_); a higher value indicates, therefore, that a higher fraction of variance is explained by individual models, rather than being shared across them. In principle, this value can take extreme values as 0, when only shared information is coded, or infinite, when only orthogonal information is represented. To evaluate whether orthogonality varies from posterior to anterior visual areas, we tested whether a linear trend between the Y coordinate and average orthogonality within XZ-slices was present by searching for abrupt changes in the slope, as high as 50% of the maximum value. As we found no significant changes, the strength of the linear dependency between orthogonality and the posterior-to-anterior direction was calculated using the Spearman’s correlation (Figure 6A) and significance was then computed with a parametric test.

**Figure 4.**
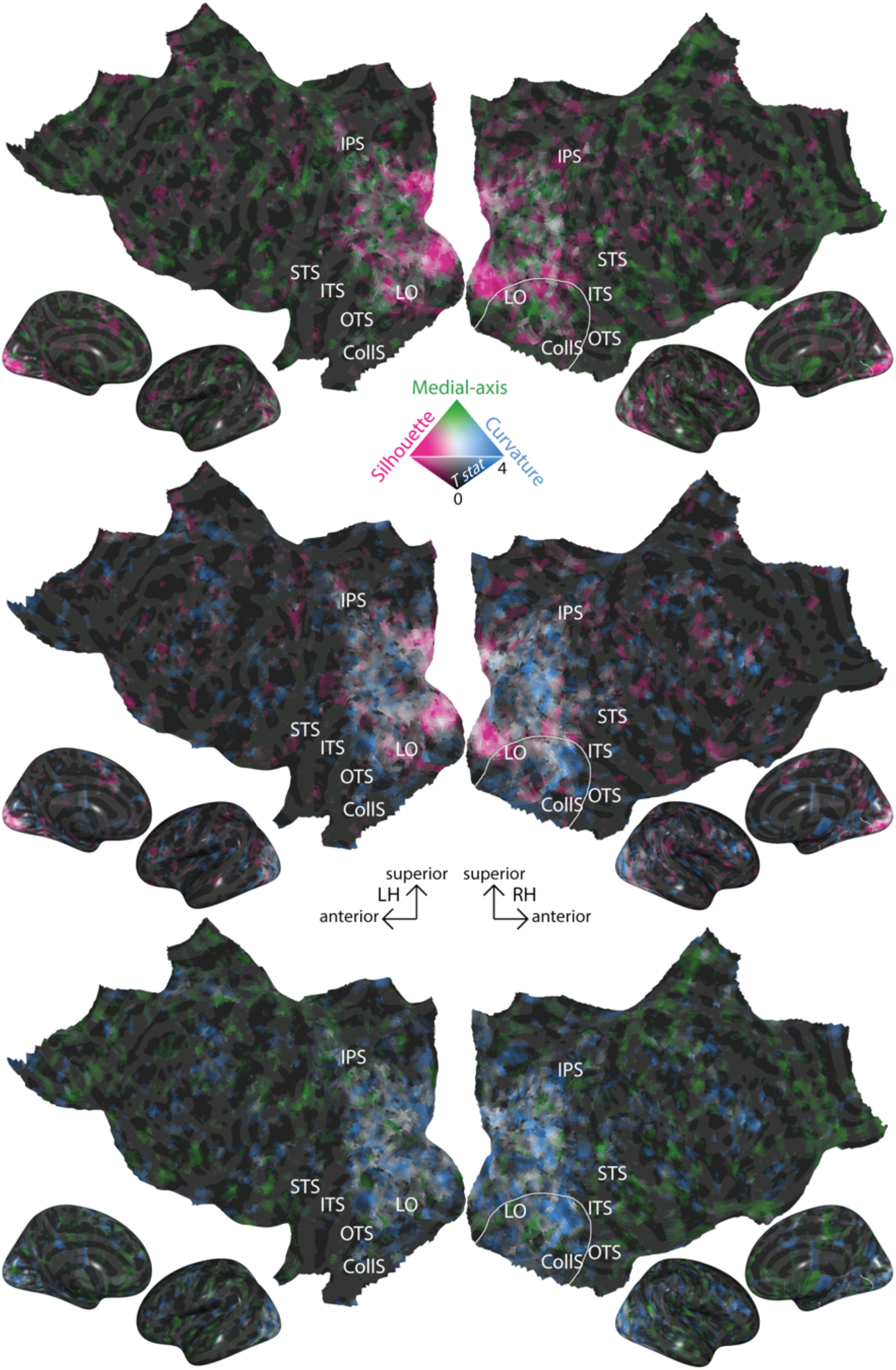
Coding of orthogonal shape components overlap in the human visual cortex. Pairwise comparisons between group-level unthresholded T-maps of orthogonal shape components show that several regions encode more than a single orthogonal description. Colored voxels have high T-value in a single model. Silhouette is represented in red, medial-axis in green and curvature in blue. The overlap between two orthogonal representations is indicated by white voxels, while brightness represents the value of the T-statistic in each voxel (i.e. gray and black voxels have low T-value in both models). White lines enclose right OTC, where all three shape models are significant.

**Figure 5.**
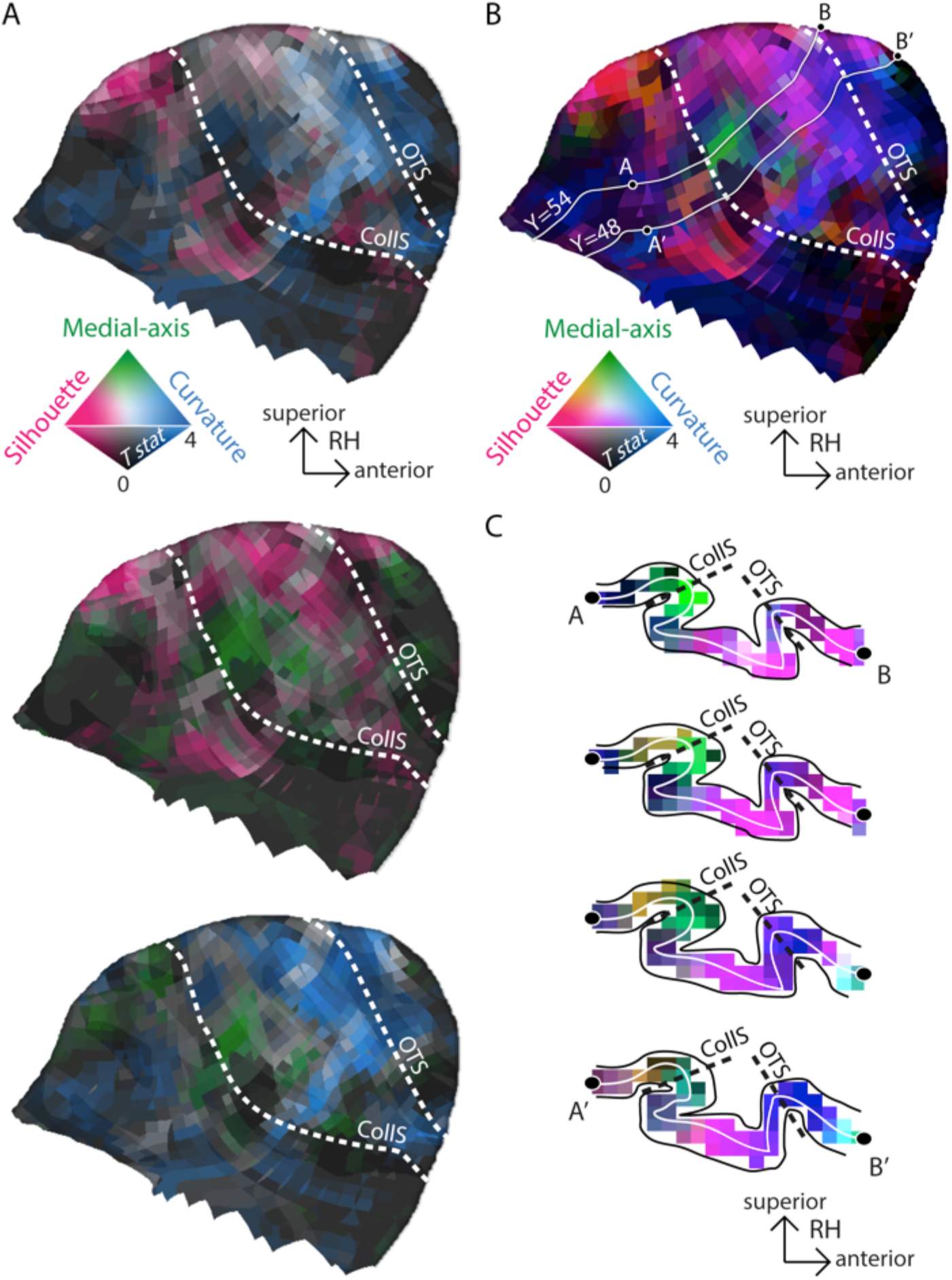
Topographic organization of object shape in right OTC. A) Pairwise comparisons between group-level T-maps of orthogonal shape components in right OTC. Colored voxels have high T-value in a single model. Silhouette is represented in red, medial-axis in green and curvature in blue. The overlap between two orthogonal representations is indicated by white voxels, while brightness represents the value of the T-statistic in each voxel (i.e. gray and black voxels have low T-value in both models). B) Overlap between the three group-level T-maps of orthogonal shape components in right OTC. Silhouette is represented in red, medial-axis in green and curvature in blue. The overlap between two orthogonal representations is indicated by intermediate colors: pink for silhouette and curvature, orange for silhouette and medial-axis, cyan for medial-axis and curvature. Brightness represents the value of the T-statistic in each voxel (i.e. gray and black voxels have low T-value in all models). The solid lines show the borders of the four slices depicted in (C): these are arranged from more posterior (top) to more anterior (bottom) and are enclosed between points (A-B and A’-B’) referring to their medial-to-lateral extent.

**Figure 6.**
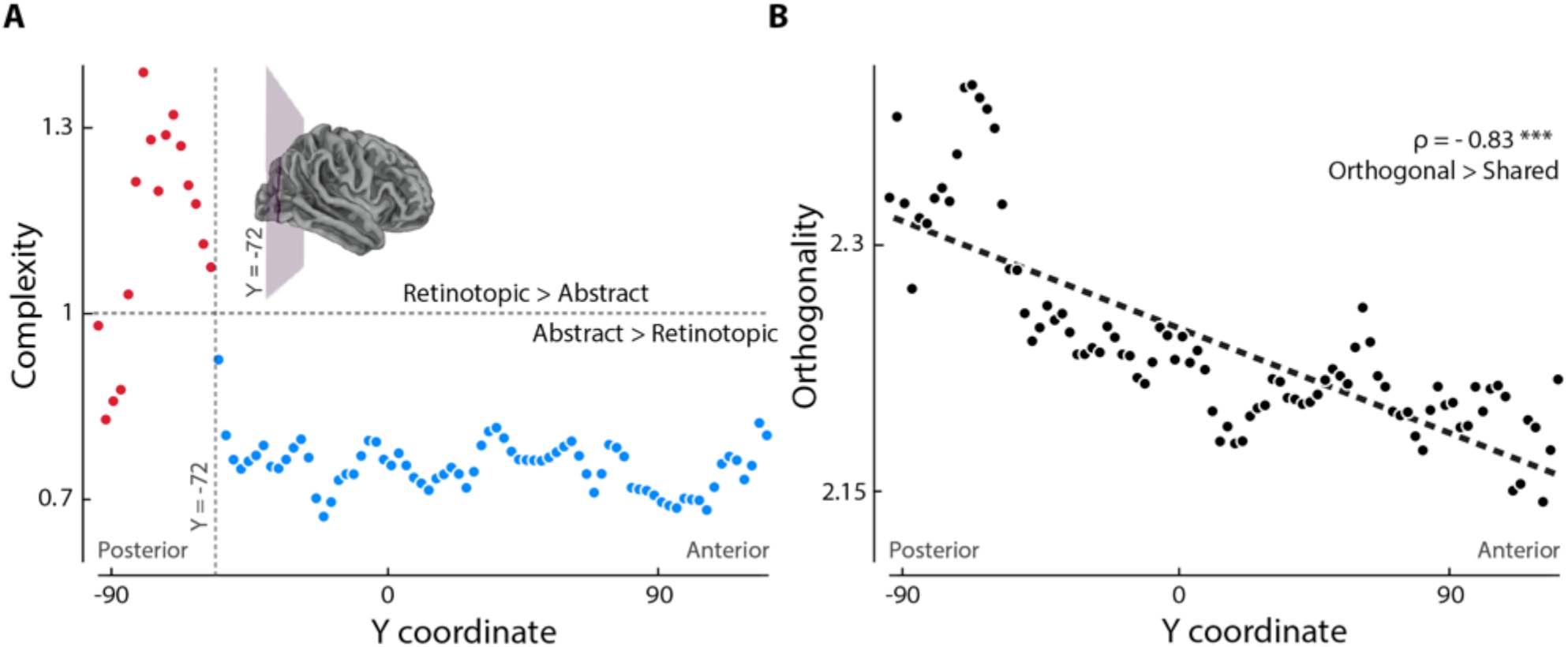
The link between object features shapes the human visual hierarchy. A) The ratio between the explained variance of low-level and abstract features (i.e. complexity) along the visual hierarchy reveals an abrupt shift. Values higher than one (horizontal dashed line) indicate that brain representations are better accounted for by retinotopic information while values smaller than one that abstract representation is more relevant. The vertical dashed line represents the point where mean and slope (dashed black lines) present an abrupt change. B) The ratio between the variance explained by the orthogonal components of the five models and the sum of the shared components between the five models (i.e. orthogonality) linearly decreases along the visual hierarchy (ρ = -0.83, ***: p < 0.001, parametric test). Values higher than one indicate that brain representations are better explained by orthogonal components of variance.

Following (Vernon et al., 2016), two different groups of features were identified: low-level representations, sensitive to retinotopic information, and abstract representations, that are independent of the extent of retinotopic cortex stimulated. Inked-area and silhouette were labeled as low-level models, and medial-axis, curvature and identity as abstract ones. Then, complexity was measured by the ratio between variance uniquely explained either by low-level or abstract models: Complexity = (R^2^ _silhouette_ + R^2^ _inked area_) / (R^2^ _medial-axis_ + R^2^ _curvature_ + R^2^ _identity_). Thus, within each searchlight, group-averaged (between subjects) sum of variance explained uniquely by the low-level models was divided by the group-averaged sum of variance explained by the abstract ones. Linearity was tested as for the orthogonality index.

Surface plots in Figures 4 and 5 were produced with the *Pycortex* toolbox for Python (Gao et al. 2015). Second-level analyses were performed using custom-made code written in MATLAB (MathWorks Inc.). Custom Matlab code is available on GitHub: https://github.com/PPthe2nd/ShapeVar. Preprocessed data for all subjects are publicly available on Zenodo: [link will be available upon acceptance]. Access to the data is possible upon submission of a Data User Agreement, available along with supplementary information on Zenodo: https://doi.org/10.5281/zenodo.4003861.

## Results

As expected from both theoretical and experimental investigations on this topic (Kay, 2011; Bracci and Op de Beeck, 2016; Papale et al., 2019), the combination of our five models and our stimulus set reveals moderate-to-high degrees of collinearity (Figure 2E). Consequently, to account for multicollinearity before considering the significance of the association of each model with brain representations, the variance partitioning analysis (Lescroart et al., 2015) and a searchlight procedure (whole brain, 6mm radius: Kriegeskorte et al. 2006) were combined to identify group-level clusters significantly explained by three shape models independently from competing representations (Figure 2D).

### The human visual cortex encodes multiple orthogonal shape representations

Group-level results show both distinct and overlapping clusters of shape selectivity in OTC, also extending to posterior dorsal regions (p < 0.05 one-tailed, TFCE corrected). The silhouette model (Figure 3, in red) shows a significant association with brain representations along the Calcarine sulcus (CalcS), the occipitotemporal sulcus (OTS), the right collateral sulcus (CollS), the right inferior temporal sulcus (ITS), the right fusiform gyrus (FusG), the cuneus (Cun) and in posterior portions of the middle temporal gyrus (pMTG) and intraparietal sulcus (pIPS). The medial-axis (Figure 3, in green) explains a significant portion of unique variance in the right lateral occipital area (LO) only. Finally, curvature (Figure 3, in blue) significantly explains fMRI representational geometries in the left lingual gyrus (LinG), in the bilateral FusG, along bilateral OTS and ITS, along the right CollS, in the right MTG, bilaterally in the Cun and along the right IPS. The significant clusters for the low- and high-level models are also represented in Figure 3.

As all orthogonal components of our tested models indicate the presence of at least a significant cluster of selectivity, shape representation does not rely on a single feature, but on a multi-dimensional coding scheme.

### Selectivity to orthogonal shape representations coexist in the same cortical regions

We further explored the overlap between the selectivity to orthogonal shape representations. Figure 4 depicts the pairwise comparisons between the three shape models in our study. Qualitatively, a stronger overlap is observed in LO for medial-axis and curvature, and in IT, right FusG, Cun, right pMTG and right pIPS for silhouette and curvature. Thus, those brain regions encode multiple shape features, independently from the shared variance between them. Also, both the effect sizes and spatial patterns are consistent across subjects (Figures S1-5).

### Topographic organization of object shape in right OTC

Of note, all three models are significant only within right OTC (enclosed by a white line in Figure 4). Figure 5 depicts right OTC in isolation with a greater detail: when combining the three models (Figure 5B), a topographic organization emerges. Silhouette coding is medial with respect to the CollS, encompassing the LinG and parahippocampal gyrus (PHG, red voxels in Figure 5B). Proceeding laterally, the silhouette and medial-axis coexist in the fundus of the CollS (orange voxels in Figure 5B), while the medial-axis extends also to the FusG (green voxels in Figure 5B). Finally silhouette and curvature are both encoded medial to the OTS, with curvature being encoded also in the fundus of the OTS.

These qualitative observations return a complex picture on shape coding in the human brain. However, some general considerations can be made by looking at the interactions between features.

### Coding of orthogonal object representations decreases from posterior to anterior regions

In a previous study, Vernon *et al*. (2016) explored the relationship between retinotopic and more abstract object representations, including contour curvature. They defined two orthogonal components enclosing low-level and higher-level, complex features, and described a shift between retinotopic and more abstract features in LO. Given the pattern of results from the previous analysis, we further explored what has been observed by Vernon et al. (2016) and also tested the relative weight of orthogonal and shared components. Indeed, the tuning to increasingly complex features is considered a cornerstone of hierarchical object processing (Riesenhuber and Poggio 2000). However, it has been proposed that interaction between features may play a pivotal role in evolving reliable selectivity in the brain (Benjamin et al. 2019). Thus, we hypothesized that shared information should become more relevant along the visual hierarchy, moving from posterior to anterior brain regions.

We defined two independent components, one for the low-level features and one for the abstract ones, as in Vernon et al (2016). The first comprised the orthogonal variance of silhouette and inked-area, since both are linked to the local retinotopic arrangement and to the extent of retinotopic cortex stimulated. The second includes the orthogonal variance of medial-axis, curvature (both insensitive to differences in object orientation and size) and object identity models. The ratio between the explained variance of low-level and abstract features (i.e., complexity) in the posterior-to-anterior axis was computed by averaging the values in the XZ plane: values higher than one indicate that brain representations are better accounted for by retinotopic information, while values smaller than one indicate that abstract representations are more relevant. When looking at the slope of complexity along the posterior-to-anterior axis, we observed an abrupt shift (higher than 50% of the maximum) from retinotopic to abstract features around Y_MNI_ = -72 (Figure 6A). Of note, the shift occurs at the limit between occipital and temporal or parietal cortex. As a matter of fact, previous studies (Haxby et al. 2001; Rice et al. 2014) on ventral temporal cortex selectivity constrained their analysis between Y_MNI_ = -70 and Y_MNI_ = -20 (but see: Grill-Spector and Weiner 2014 for a different definition based on anatomical landmarks).

Then, we looked at the ratio between orthogonal and shared variance components (i.e., orthogonality) in the posterior-to-anterior axis. The variance explained by the orthogonal components of the five models was first summed, and then divided by the sum of the shared components between the five models. Here, values higher than one indicate that brain representations are better explained by orthogonal components of variance. Orthogonality linearly decreases along the posterior-to-anterior axis, without shifts (ρ = -0.83, p < 0.001, parametric test; Figure 6B). Thus, while orthogonal information is always more represented than shared variance (min = 2.15), it becomes less relevant proceeding along the visual hierarchy, and neither ventral nor dorsal streams alone are responsible for the overall effect (Figure S6).

Of note, both these ratios abstract away from magnitude of response and goodness of fit *per se*. Instead, they reveal the relative contribution of different features to different brain areas beyond their overall performance.

## Discussion

In the present study, we found that object shape is not represented by a single feature but is encoded by multiple representations (i.e., silhouette, medial-axis and curvature) that uniquely contribute to object processing in the human visual cortex (Figures 3-5). Moreover, we showed that the brain encodes orthogonal object representations in a topographic fashion: the early visual cortex is biased towards unique components of variance, while shared representations become progressively more relevant in more anterior regions (Figure 6).

### Shape coding is multidimensional

In line with previous studies, we found that object silhouette is mainly encoded in early visual areas (Bracci and Op de Beeck 2016; Kaiser et al. 2016; Khaligh-Razavi and Kriegeskorte 2014; Proklova et al. 2016). This result can be explained by top-down figure-dependent mechanisms that modulate V1 activity both in monkeys (Poort et al. 2016; Self et al. 2019) and humans (Kok and de Lange 2014; Muckli et al. 2015), and enhances the processing of object-related information in early visual areas also during natural vision (Papale et al. 2018). However, another possibility may be that the silhouette model better captures the object physical appearance (Kubilius et al. 2016).

Instead, the variance component unique to the medial-axis model – which is the most transformation-resistant shape description (Yang et al. 2008) – was significant in a smaller extent of cortex comprising only a subset of voxels in right LO (Figure 3, middle in green). This can be due to a higher spatial inter-subject variability of this representation that has been already observed by Leeds *et al*. (2013), or to a higher collinearity with the low- and high-level models we employed (Figure 2D) that prevents from disentangling its contribution from competing representations. Nonetheless, our result fits previous evidence of medial-axis coding in monkey IT (Hung et al. 2012; putative homologue of human LO), human LO (while also controlling for low-level properties: Ayzenberg et al. 2019b) and is consistent with our previous MEG study showing that medial-axis processing is limited to a small cluster of right posterior sensors, when controlling for collinearity with low-level and categorical representations (Papale et al. 2019).

Finally, IT (Kayaert et al. 2005b; Yue et al. 2014), LO (Vernon et al. 2016) and FusG (Caldara et al. 2006) were bilaterally tuned to contour curvature (Figure 3, bottom in blue), in accordance with previous neuroimaging investigations. Actually, LO has a pivotal role in object processing (Grill-Spector et al. 2001; Grill-Spector et al. 1999; Kourtzi and Kanwisher 2001), as IT in monkeys (Brincat and Connor 2004; Desimone et al. 1984; Kayaert et al. 2005a; Op de Beeck et al. 2001; Tanaka 2003; Zoccolan et al. 2007). In addition, while we focus our discussion on the ventral stream, we also observed few significant clusters in dorsal visual regions (R pIPS; see Figure 3), both for curvature and silhouette, which confirm previous observations (Freud et al. 2017).

Our result favors the hypothesis of a key role of right OTC in coding object shape. In line with our view, recent neuropsychological evidence found visual agnosia in a subject with a cortical lesion circumscribed to right OTC. Moreover, this focal right OTC lesion affected shape selectivity across the whole visual cortex, leading to large-scale alterations that were stable over time (Freud and Behrmann 2020).

Overall, the evidence that all the tested dimensions independently contribute to shape representation in the human visual cortex favors the hypothesis of a multi-dimensional coding of object shape (Silson et al. 2016; Silson et al. 2013), similarly to what is observed for texture processing (Okazawa et al. 2015; Ziemba et al. 2016). In line with this, a recent study showed that behavioral shape similarity could be modeled with no less than 109 different dimensions (Morgenstern et al. 2020).

### Coding of shared information increases along the visual hierarchy

Long *et al*. (2018) suggested that mid-level computations, covarying with high-level semantic processing (including curvature extraction), control the organization of OTC. In the present study, however, we observed overlapping selectivity to orthogonal features in LO (medial-axis and curvature), IT, right FusG, Cun, right pMTG and right pIPS (silhouette and curvature). Since we controlled for collinearity between models, this result could not be merely ascribed to the variance shared by those features. Here, we also observed that coding of shared descriptions in OTC is topographically arranged and its relevance linearly increases from posterior to anterior regions (Figure 6). This observation, consistent with the core finding of Long *et al*. (2018), suggests that the hierarchy of visual processing is not only shaped by specificity to increasingly complex features, but also by a higher selectivity to shared representations.

This observation complements what has been already observed on the two extremes of the ventral visual pathway: V1 and IT. Representations in V1 are over-complete relative to the retinal input (Olshausen and Field 1996; Vinje and Gallant 2000). In addition, inhibitory interactions in V1 are specifically targeted at neurons with similar tuning properties (Chettih and Harvey 2019). Both these factors increase V1 representational capacity and may ultimately lead to a higher selectivity to orthogonal features, as we observed in posterior regions. On the other hand, higher sensitivity to shared information in more anterior areas may be produced by populations of neurons that are not tuned to a specific property but that encode multiple dimensions at once. Indeed, shared featural selectivity has been proposed as the mechanism responsible to achieve dimensionality reduction of the sensory input in IT (Lehky et al. 2014), where both neural density and surface are much lower than in V1 (Cahalane et al. 2012; Van Essen et al. 1992). In line with this, the highest-dimensional among our three shape models (i.e., silhouette) is also represented in posterior regions (Figure 3). Relatedly, the interaction between multiple features is thought to represent the optimal solution to increase the sensitivity to their mutual changes: in this view, instead of having few neurons encoding a single feature each, it may be preferable to have most of the neurons encoding multiple features at once (Benjamin et al. 2019). It has been also suggested that interactions between features are responsible for the poor reliability of tuning curves in predicting brain responses in natural vision (Benjamin et al. 2019).

Thus, what can be concluded on the nature of object processing? On one hand, we observed an abrupt shift from retinotopic to abstract representations moving anteriorly across the brain (Figure 6A). However, this shift is relative: though less relevant, orthogonal retinotopic information spreads also to OTC, explaining a significant portion of its variance, in line with previous work and suggesting a link between low-level and object selectivity (Rajimehr et al. 2011; Rice et al. 2014). On the other hand, we found a linear dependency between the posterior-to-anterior axis and the variance explained by shared information (Figure 6B), in line with previous research showing an increase in coding category-orthogonal information from V4 to IT in non-human primates (Hong et al. 2016). As stated earlier, this property describes the linear cascade of computations in the visual hierarchy better than complexity: optimizing the coding of shared variance between behaviorally relevant features may represent a key factor in shaping the architecture of our visual cortex and achieving reliable, view-point invariant object representations. In this light, the next step should be to move from modeling representational geometries to more direct modulations of brain responses, so to control also for nonlinear interactions between features (Benjamin et al. 2019).

### Limits and conclusions

It should be noted that due to the low fMRI temporal resolution, we cannot resolve which mechanisms support the different tuning for shared representations. Moreover, while the selected models capture visual transformations, many alternative descriptions exist (e.g., Khaligh-Razavi and Kriegeskorte 2014). In addition, the searchlight procedure introduces some unavoidable imprecision in localization (but see: Lettieri et al. 2019 for an analytical exploration of this issue), thus further studies using the same shape descriptions might found slightly different locations as those found in this study. Overall, however, our results hint at the existence of a multi-dimensional coding of object shape, and reveal that selectivity for shared object representations are topographically arranged and increases along the visual hierarchy. Future experiments will identify how different tasks (e.g., determining object similarity vs. extracting affordances), and alternative descriptions impact on the observed patterns of selectivity. Finally, we described what qualitatively appears to be a shape coding topography in right OTC: further research would be necessary to understand how strong is the link between structural and functional organizations.

## Acknowledgments

This work has been supported by the Italian Ministry of Education, University and Research grants PRIN 2015WXAXJF and 2015AR52F9.

## Author contributions

Pa.P., A.L. G.H., L.C. and E.R. conceived the study. A.L., G.H. and L.C. performed experiments. Pa.P., and A.L. analyzed the data. All the authors discussed the results and wrote the manuscript.

